# Chronic pain increases sensitivity to pain-induced reinstatement of ethanol seeking in male mice

**DOI:** 10.1101/2025.04.10.648244

**Authors:** Mitchell A Nothem, Christine M Side, Simon C Tran, Anaahat Brar, Lauren A Buck, Jacqueline M Barker

## Abstract

Alcohol use disorder (AUD) and chronic pain are complex and debilitating conditions that are highly comorbid. Greater than 50% of individuals with AUD have chronic pain. Clinical data suggest that people with chronic pain are more likely to report using alcohol to manage chronic pain, and that magnitude of pain is correlated with relapse probability after a period of abstinence. These data led to the hypothesis that pain can drive ethanol seeking and reinstatement in a rodent model of chronic neuropathic pain. A conditioned place preference (CPP) paradigm was used to model ethanol seeking in male C57BL6J mice with a spared nerve injury (SNI). Mice were conditioned with doses of ethanol previously found to reverse pain behavior (0.5g/kg). Mice with and without SNI showed similar magnitudes of ethanol CPP and rates of extinction. To investigate pain-induced relapse-related behavior, mice underwent reinstatement testing following painful mechanical stimulation which was delivered at either a “moderate” or “high” intensity immediately prior to return to the CPP apparatus. “Moderate” painful hindpaw stimulation reinstated ethanol seeking behavior in SNI-injured, but not sham, mice, while “high” intensity stimulation reinstated ethanol seeking in mice regardless of injury status. These data suggest that males in chronic pain are more susceptible reinstatement of ethanol seeking following a painful experience.

## Introduction

Alcohol use disorder (AUD) and chronic pain are highly comorbid conditions that significantly impair quality of life measures and impose substantial social and economic burdens to society (Andrew et al., 2014; Axley et al., 2019). Over half of individuals with AUD also suffer from chronic pain (Boissoneault et al., 2019). This co-occurring AUD and chronic pain are associated with anxiety, depression, opioid use, and increased substance use disorders (Boissoneault et al., 2019). People in chronic pain frequently report pain to be a primary driver of their alcohol use and upwards of one -third of binge drinkers consume alcohol to self-medicate pain (Alford et al., 2016). Further, chronic pain is associated with greater risk of relapse (Boissoneault et al., 2019; McDermott et al., 2018; Von Korff et al., 2005). This is consistent with modulatory effects of alcohol on sensory thresholds and pain tolerance that are comparable to traditional analgesics (Cutter & O’farrell, 1987; Horn-Hofmann et al., 2015; Perrino et al., 2008; Wolff et al., 1940; Woodrow & Eltherington, 1988). However, alcohol’s analgesic properties are subject to the development of tolerance and can lead to increased consumption, which then can worsen pain outcomes (Brennan et al., 2005; Egli et al., 2012) Thus, the relationship between chronic pain and AUD is bidirectional and difficult to investigate in the clinical setting.

Preclinical models can be used to better understand the relationship between chronic pain and AUD and mechanisms that drive AUD-related behaviors. We and others have demonstrated that ethanol is able to reduce pain-like behaviors in a variety of animal models (Dundee et al.,1969.; Horn-Hofmann et al., 2019; King et al., 2013; Niesters et al., 2014; Nothem et al., 2023). For example, we previously reported that in the spared nerve injury (SNI) model of chronic neuropathic pain, ethanol reduces mechanical allodynia and reverses shifts in dynamic weight bearing (DWB) across all tested doses in male mice (Nothem et al., 2023). The bidirectionality of this relationship is further modeled in rodents, as the presence of chronic pain increases ethanol seeking and consumption in mice (Butler et al., 2017; Yu et al., 2019).

A defining feature of AUD is relapse to alcohol use, even after extended abstinence. As chronic pain is associated with greater risk for relapse, we investigated the impact of a history of chronic pain on a novel model of pain-induced relapse-related behavior. As we previously reported that the ability of ethanol to reverse pain-related behavior in females with spared nerve injury was modest (Nothem et al., 2023), we investigated conditioned place preference (CPP) for a low dose of ethanol that reversed pain-related behavior only in males with SNI (Decosterd & Woolf, 2000; Erichsen & Blackburn-Munro, 2002; Guida et al., 2020). After ethanol CPP conditioning and extinction, we assessed whether painful stimulation would reinstate a place preference. Our findings demonstrate similar ethanol CPP in male mice with and without SNI, while mice with chronic neuropathic pain exhibited greater sensitivity to pain-induced reinstatement. These findings suggest that a history of chronic neuropathic pain enhances the ability of painful stimuli to elicit relapse-related behavior and may provide a valuable model to develop strategies to suppress relapse to alcohol use in individuals with a common comorbidity.

## Methods

### Animals

Adult male C57Bl/6J mice were used (Jackson Laboratories). Mice were 9 weeks old and between 18-23 g at the beginning of the experiments. All animals were housed in a temperature and humidity-controlled environment on a standard 12h:12h light-dark cycle and provided food and water *ad libitum* for the duration of the experiment. All behavioral experiments occurred within the light cycle. Mice were handled and allowed to acclimate to the colony room for one week prior to beginning the baseline behavior experiments. In total, 26 animals were used in these studies (Sham: 13; SNI: 13). All procedures were carried out under a protocol approved by the Institutional Animal Care and Use Committee of Drexel University.

### Spared nerve injury (SNI) surgery

Mice underwent SNI surgery (Decosterd & Woolf, 2000) at approximately 11 weeks of age. Mice were anesthetized by inhaled isoflurane. An incision was made on the left hindleg, and the sciatic nerve was exposed. The common peroneal and tibial nerves were ligated with 6-0 silk suture and cut distally from the ligation. The skin was closed using size 7 wound clips (Reflex). The sural nerve was left intact. Sham mice underwent a similar procedure in which the branches of the sciatic nerve were exposed but not ligated or cut. Animals recovered for 2 weeks for the development of hypersensitivity to occur and for changes in weight bearing to stabilize prior to behavioral testing (**Fig 1a)** (Mogil et al., 2010). Mice were excluded if they did not develop mechanical allodynia after SNI surgery defined as having a paw withdrawal threshold greater than 0.25g at the post-surgery timepoint. One animal was excluded due to this criterion.

**Figure 1.**
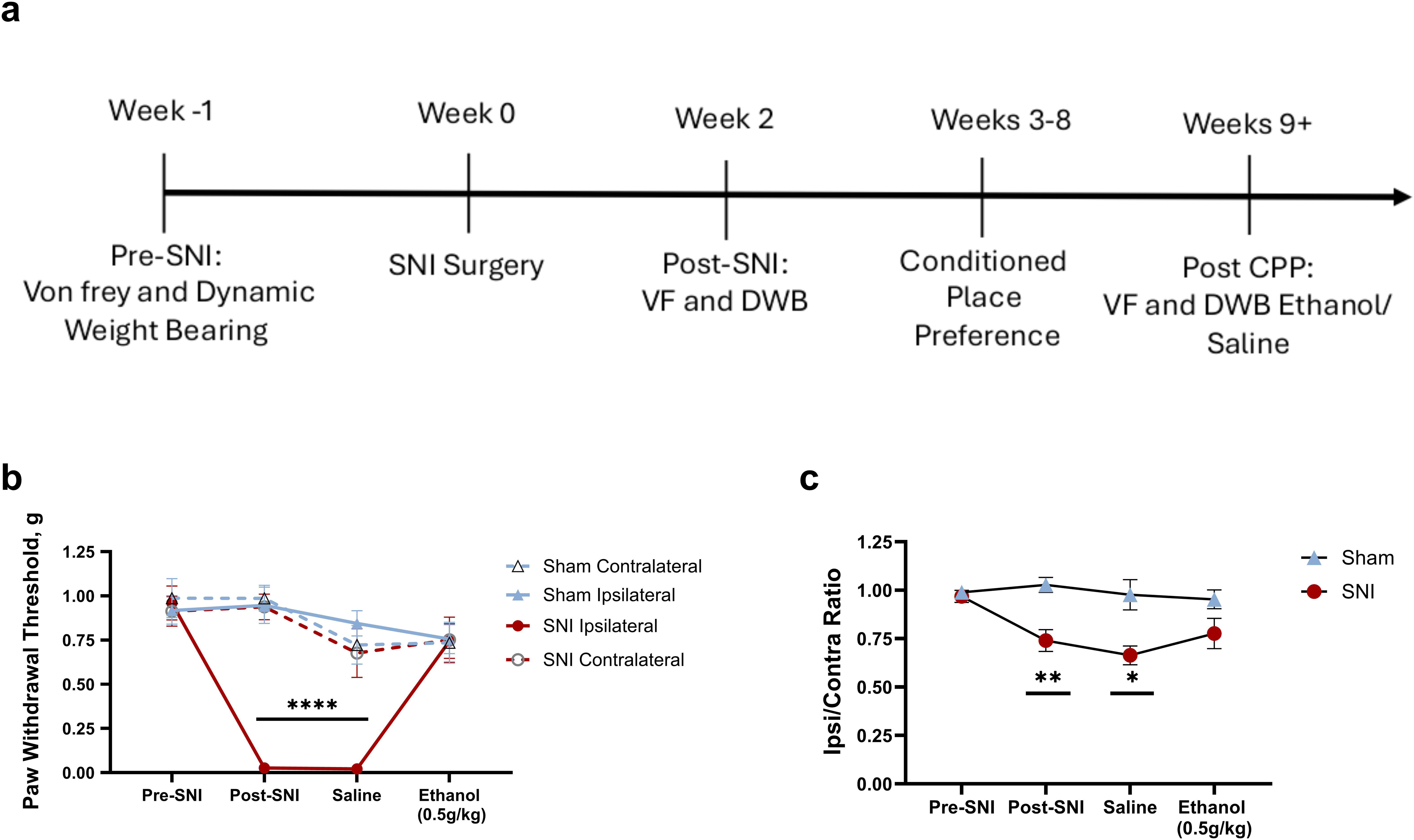
Experimental timeline and effect of SNI and ethanol on pain-related behavior. **(a)** Timeline of SNI and surgery followed by behavioral testing. **(b)** Paw withdrawal thresholds measured using the von Frey assay were significantly reduced in the ipsilateral hindpaw of SNI mice compared to sham controls before and after CPP testing. This was reversed by acute injection of 0.5g/kg ethanol. **(c)** Dynamic weight bearing analysis showed decreased weight distribution on the injured hindlimb in SNI mice compared to shams before and after CPP. Treatment with 0.5g/kg normalized weight bearing in SNI mice. Data are expressed as mean ± SEM. *p < 0.05 vs. sham; **p < 0.01 vs. sham; ***p < 0.001 vs. sham.

### Von Frey

To assess mechanical hypersensitivity, mice were stimulated with a set of von Frey fibers using the up-down method, beginning with a 0.16g fiber (Stoelting Touch Test, Chicago, IL) and with 2.56 g as the upper limit (Chaplan et al., 1994; Dixon, 1980). Animals were tested prior to surgery to establish baseline sensitivity and then two weeks after surgery, which was prior to ethanol conditioned place preference (CPP) training and testing. After completing the CPP paradigm, mice were tested 10 minutes after with an injection of saline or ethanol. The dose of ethanol was identical to the dose used in CPP conditioning (0.5 g/kg).

### Dynamic weight bearing (DWB)

To assess the effect of SNI surgery and acute ethanol administration on DWB, behavior was recorded using a DWB apparatus (Dynamic Weight Bearing 2.0, Bioseb, France) prior to and 2 weeks after SNI surgery (Nothem et al., 2023). After completing the CPP paradigm, mice were injected with saline or ethanol and tested 10 minutes after injection using the same 0.5 g/kg dose used in CPP conditioning. Throughout testing, mice were acclimated for 5 minutes prior to 5 min of recorded behavioral data. During acclimation and recordings, mice were able to freely rear and explore the chamber, while a grid of floor force sensors measured the weight placed on each hindpaw and an overhead camera recorded the animal’s position and posture. Paw detection settings were defined with a central pixel sensitivity threshold of 0.8 g and an adjacent pixel threshold of 0.2 g, allowing for reliable detection of hindpaws. A minimum of 60 seconds of video was validated using a combination of Bioseb’s automatic validation (Level 1 and 2) and manual validation to ensure that paws were accurately identified. Animals were excluded from analysis if 60 seconds of data could not be validated from the recordings due to technical issues. At the saline timepoint 2 shams, and 3 SNI’s were excluded and at the ethanol time point 3 SNI animals were excluded due to lack of greater than 60 seconds of validation.

### Conditioned place preference

The CPP apparatus consisted of two distinct compartments with different visual and tactile cues (black walls with bar flooring and white walls grid floors) connected by a neutral gray chamber with smooth floors (Med Associates, St. Albans, VT). The paradigm was conducted over 8 days and included three phases: pre-conditioning, conditioning, and CPP test. Entry into chambers and locomotor activity were automatically detected using photobeams.

During preconditioning, mice were allowed to explore the entire CPP apparatus for 20 minutes. The time spent in each compartment was recorded to ensure no group (SNI vs Sham) differences were present in initial preference for either compartment. Mice underwent six days of conditioning sessions, with alternating exposure to 0.5 g/kg ethanol and saline via i.p. injection. During conditioning days animals were injected with either ethanol or saline and immediately confined to one compartment for 20 minutes. The assignment of ethanol and saline to specific chambers and order of injection was counterbalanced across mice to control for compartment bias. During the post conditioning CPP test, mice were placed into the neutral chamber with the doors opened and were allowed to explore the entire CPP apparatus for 20 minutes. The time spent in each compartment was recorded. A preference score was calculated as the difference in time spent in the ethanol-paired compartment during the post-conditioning session compared to the pre-conditioning session.

### Extinction Training

After the post-conditioning test and pain-induced reinstatement tests, mice underwent extinction training where they were placed into the neutral gray chamber and allowed to explore each chamber for 20 minutes. Extinction training continued daily until 80% of mice reduced their preference for the ethanol-paired chamber, mice were excluded from reinstatement tests if time spent in the ethanol-paired chamber was 200 seconds greater than the saline-paired chamber on the last extinction day. A maximum of 30 days of extinction training sessions were used as a cutoff point, none of the groups of mice reached 30 days of extinction training sessions.

### Pain-induced reinstatement tests

On moderate reinstatement test day, animals underwent von Frey testing to calculate a 50% paw withdrawal threshold for each animal (PWT). After PWT’s were calculated animals were stimulated 10 times on the lateral aspect of the plantar surface of the ipsilateral hindpaw relative to injury with a von Frey fiber that delivered a force corresponding to the ∼50% PWT and number of hindpaw responses were recorded. Reinstatement stimulations occurred within 1-2 minutes, depending on animal guarding and rearing behaviors. Immediately after stimulations, animals were placed in the neutral chamber and allowed to freely explore all 3 chambers of the CPP apparatus for 20 minutes. Time spent in each chamber was recorded.

Procedures for ‘moderate’ stimulation reinstatement versus ‘high’ stimulation reinstatement were identical with the exception being the amount of force used to deliver the 10 stimulations prior to the reinstatement test. ‘Moderate’ stimulation reinstatement was defined as using the von Frey fiber closest to the calculated 50% withdrawal threshold, while high stimulation reinstatement test used 2 fibers above the 50% PWT, theoretically equivalent to a ∼90% PWT. Reinstatement tests were counterbalanced such that 50% of animals received moderate stimulation reinstatement first while the other half of the animals received the high stimulation reinstatement first to account for repeated testing.

### Statistics

All statistical analysis was conducted using Prism GraphPad. Analyses consisted of between-subjects or repeated measures (rm) ANOVA, and one- or two-sample t tests as specified in the results. In the case of missing values in repeated measures tests, a mixed-effects design was used. Significant interactions were followed by Sidak’s post hoc comparisons.

## Results

### Spared nerve injury induces mechanical allodynia and shifts in weight bearing

To confirm mice developed mechanical allodynia after SNI, basal paw withdrawal thresholds on ipsilateral and contralateral hindpaws were measured prior to and after SNI surgery using the von Frey and dynamic weight bearing assays. After conditioned place preference (CPP) behavior, animals were given either an i.p. injection of saline or 0.5g/kg of ethanol and underwent an additional round of testing to confirm the effect of ethanol on mechanical allodynia and weight bearing. For the von Frey assay, a 3-way rmANOVA revealed a significant 3-way injury x hindpaw x time interaction [F(3,172) = 7.678, p = <0.0001; **Fig. 1b**], indicating that SNI surgery induced changes in mechanical hypersensitivity after surgery in the injured paw. Additionally, significant two-way interactions were observed [injury x paw: F(1,172) = 17.82, p < 0.0001; time x paw: F(3,172) = 6.350, p = 0.0004; time x injury: F(3,172) = 8.145, p < 0.0001]. Main effects of time, injury, and paw were also observed [time: F(3,172) = 11.36, p < 0.0001; injury: F(3,172) = 25.44, p < 0.0001; injury: F(3,172) = 16.39, p < 0.0001]. Significant interactions were deconstructed using Sidak’s post hoc tests and revealed that SNI ipsilateral paw withdrawal thresholds were not significantly different from SNI contralateral paw (p > 0.05) or sham ipsilateral paw (p > 0.05) during the pre-SNI timepoint. However, after SNI surgery, SNI ipsilateral paw withdrawal thresholds were reduced compared to SNI contralateral paw (p < 0.0001) and sham ipsilateral paw (p < 0.0001), confirming that SNI animals developed mechanical allodynia on the injured hindpaw.

After CPP testing, paw withdrawal thresholds were tested 10 minutes after saline or ethanol injection. After saline injection, SNI ipsilateral paw withdrawal thresholds were significantly lower compared to SNI contralateral paw or sham ipsilateral paw thresholds (p’s < 0.001), indicating no effect of saline. However, after ethanol, SNI ipsilateral paw withdrawal thresholds did not differ from SNI contralateral paw or sham ipsilateral paw thresholds (p’s > 0.05), indicating that 0.5g/kg of ethanol fully reversed mechanical allodynia. Sham paw withdrawal thresholds were not affected when compared to the saline timepoint (all p’s > 0.05) suggesting that this dose of ethanol is not antinociceptive in sham animals.

The effect of SNI surgery and ethanol treatment on ipsilateral/contralateral hindpaw ratios was assessed using dynamic weight bearing (DWB), where a ratio of 1 would indicate equal weight distribution between hindpaws. A 2-way mixed effects analysis revealed a significant time x injury interaction [F(3,52) = 6.220, p = 0.0011; **Fig. 1c**]. Sidak’s post hoc tests were used to determine the effects of the interaction. Prior to SNI surgery, there was no difference between SNI and Sham groups (p = 0.9866) However, after SNI surgery, SNI animals exhibited a reduction in the ipsilateral/contralateral ratio of hindpaw weight bearing (p = 0.0021), indicating that SNI animals shifted weight towards the contralateral side and away from the ipsilateral side. This effect persisted after CPP testing, and SNI animals exhibited lower ipsilateral/contralateral ratios than sham animals after a saline injection (p = 0.0181). Consistent with our previous findings, following an injection of 0.5g/kg of ethanol, SNI and Sham groups exhibited similar ratios (p = 0.2784), indicating that ethanol reduced aberrant distribution of hindpaw weight (Nothem et al., 2023).

### SNI does not impact acquisition or extinction of an ethanol CPP in male mice

To determine the effect of SNI surgery on expression of ethanol CPP, mice underwent the experimental timeline represented in **Fig. 2a**.

**Figure 2.**
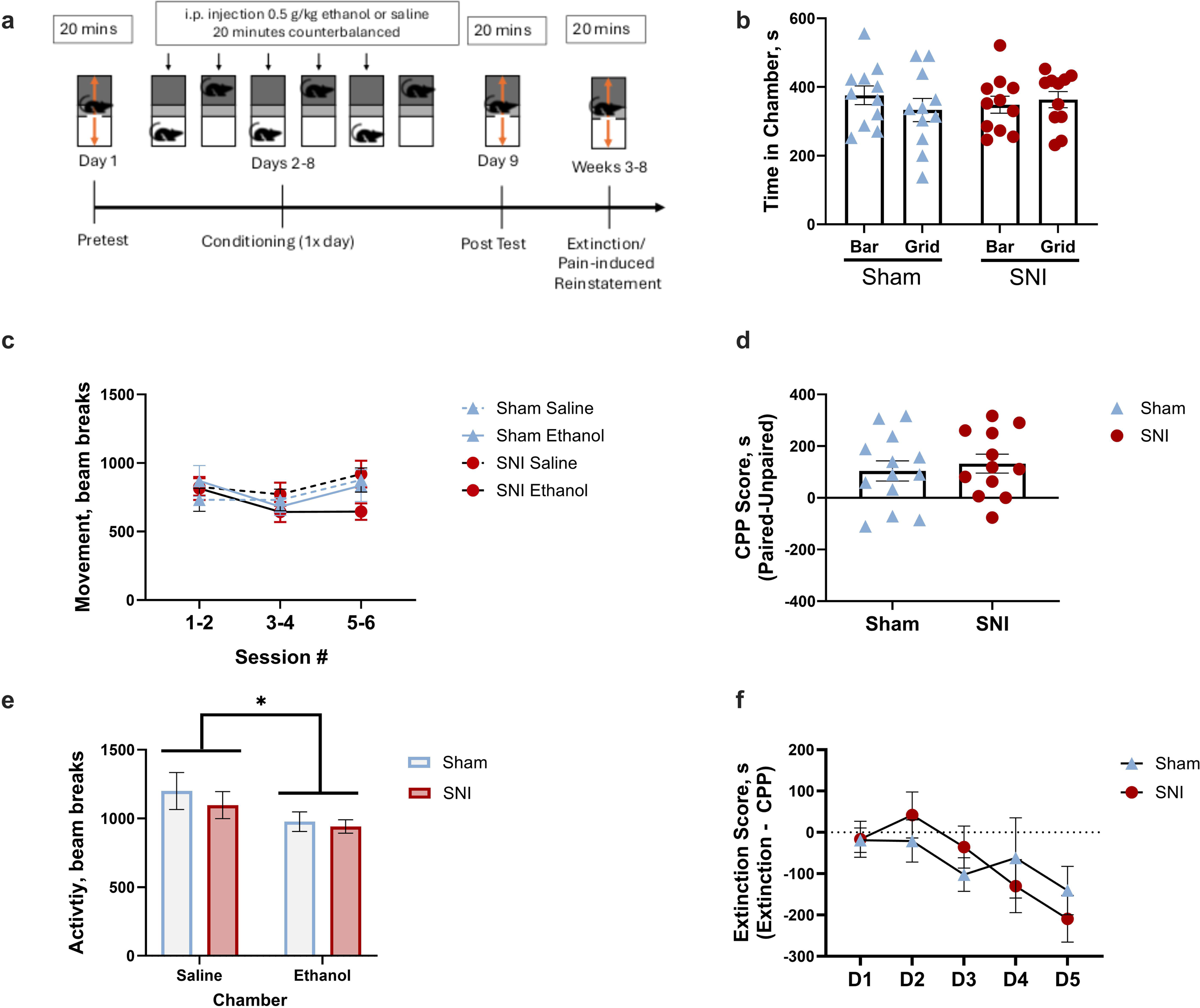
Ethanol produces conditioned place preference in SNI and sham mice. **(a)** Timeline of CPP paradigm. Mice received 3 pairings of ethanol (0.5 g/kg, i.p.) or saline in an unbiased design. **(b)** No effect of flooring was observed in either group. **(c)** SNI surgery and ethanol had no effect on movement during conditioning. **(d)** SNI and sham animals formed a CPP of similar magnitude for the ethanol-paired chamber. **(e)** Locomotor activity was reduced in the ethanol-paired chamber during CPP test. **(f)** No effect of SNI on extinction learning was observed. Data are expressed as mean ± SEM. *p < 0.05 ethanol vs saline.

The CPP chambers used in these experiments have different flooring in distinct chambers: bar floors versus grid floors. SNI mice experience mechanical hypersensitivity on the ipsilateral paw, and thus we confirmed that SNI did not impact floor type preference by assessing the amount of time spent in each chamber during the pretest. A 2-way ANOVA confirmed that there was no flooring type x injury interaction [F(1,40) = 1.08, p = 0.3029], or main effects of chamber [F(1,40) = 0.25, p = 0.6176] or injury [F(1,40) < 0.001, p = 0.9592] on time spent in each chamber prior to conditioning (**Fig. 2b**). This indicates that neither SNI nor sham animals had an initial bias for either chamber when first exposed to the CPP apparatus.

During saline and ethanol conditioning sessions, activity was measured by the total number of photobeam breaks during each session. A 3-way rmANOVA (pairing session x injury x drug) revealed that a main effect of pairing session was observed [F(1.686, 38.78), = 3.806, p = 0.0376; Greenhouse-Geisser corrected; **Fig. 2c**]. No other main effects or interactions were significant [injury x drug: F(1, 23, = 1.60), p = 0.2175; pairing session x drug: F(1.709, 39.30) = 2.251, p = 0.1257; Greenhouse-Geisser corrected; pairing session x surgery: F(2, 46) = 0.5883, p=0.5594; drug: F(1, 23) = 1.013, p = 0.3247; injury: F(1, 23) = 0.06535, p = 0.8005; pairing session: F(1.686, 38.78) = 3.806, p = 0.0376; Greenhouse-Geisser corrected]. The main effect of pairing session was deconstructed using Sidak’s post hocs but did not reach significance (all p’s > 0.05).

To determine whether SNI and sham mice developed a CPP for the ethanol-paired chamber, CPP scores were calculated by subtracting the time spent in the saline-paired chamber from the time spent in the ethanol-paired chamber. Separate one-sample t-tests were conducted for each group vs. 0. The analysis revealed that both SNI mice [t = 3.629, df = 11, p = 0.0040] and sham mice [t = 2.674, df=12, p = 0.0203) developed a significant preference for the ethanol-paired chamber (**Fig. 2d**). An unpaired t-test revealed no significant difference in CPP scores between groups [t = 0.5289, p = 0.6019].

During the CPP test, activity was measured in each chamber as the mice explored the CPP apparatus. A 2-way ANOVA (injury x chamber) revealed a significant main effect of chamber [F(1, 23) = 4.850, p = 0.0379; **Fig. 2e**], indicating less activity in the ethanol-paired chamber irrespective of injury, even though no ethanol was administered prior to the CPP test. No significant chamber x injury interaction [F(1, 23) = 0.1545, p = 0.6979] or main effect of injury [F(1, 23) = 4.850, p = 0.0379] were observed.

To determine the effect of SNI on extinction learning, SNI and sham mice underwent extinction training until CPP was significantly extinguished. A mixed effects analysis of injury x time was used to determine the effect of SNI on extinction scores and revealed a main effect of time [F(2.798, 55.96) = 5.286, p = 0.0034, Geisser-Greenhouse corrected; **Fig. 2f**] However, the analysis revealed no injury x time interaction [F(4, 80) = 2.226, p =.0734] or main effect of surgery [F(1, 22) = 0.0027, p = 0.9585]. Sidak’s post hoc analysis indicated a significant reduction on extinction day 5 vs 1 (p = 0.0110).

### Mechanical sensitivity was stable across testing

Pain-induced reinstatement testing was counterbalanced between the ‘moderate’ and ‘high’ stimulation test sessions. To confirm that repeated testing did not impact mechanical sensitivity, paw withdrawal thresholds were compared for each test session. As expected, SNI mice exhibited lower paw withdrawal thresholds than shams [F(1,20) = 207, p < 0.0001; **Fig 3a**] at the pain-induced reinstatement tests. No main effect of time [F(1,20) = 0.73, p = 0.4000] or time x injury interaction [F(1,20) = 0.53, p = 0.4729] were observed. This indicates that paw withdrawal thresholds were not different on pain-induced reinstatement day with moderate versus with high stimulation.

**Figure 3.**
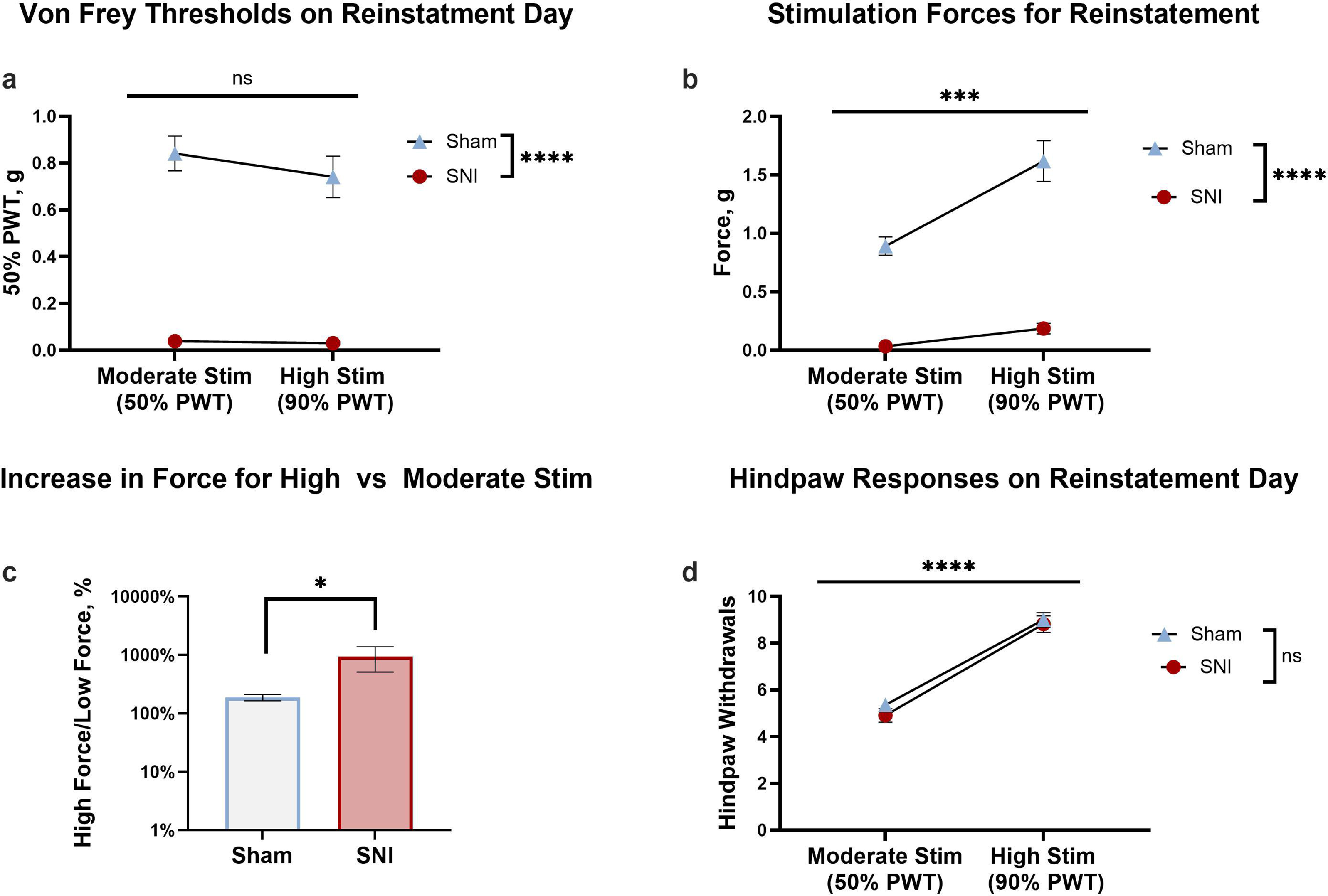
Characterization of painful stimulation in SNI and Sham animals. **(a)** While Sham and SNI animals exhibited distinct paw withdrawal thresholds, there was no difference in thresholds between test days moderate stimulation and high stimulation pain-induced reinstatement days between. **(b)** A higher force was used for high stimulation sessions than moderate stimulation sessions for Sham animals. **(c)** Proportional increase in force used to stimulate SNI animals was greater than shams between moderate and high pain-induced stimulation days**. (d)** Despite differences in force magnitude used to stimulate SNI and sham animals, behavioral responses were similar. Data are expressed as mean ± SEM. *p < 0.05 SNI vs. Sham ***p < 0.001 moderate stimulation vs. high stimulation ***p < 0.05 moderate stimulation vs. high stimulation **** p <0.0001 moderate stimulation vs high stimulation.

To compare the amount of force applied, a two-way rmANOVA indicated a significant reinstatement day x injury interaction [F(1, 20) = 9.512, p = 0.0059; **Fig. 3b**]. Sidak’s post hoc tests revealed that the amount of force applied to the ipsilateral paw of sham animals on high pain-reinstatement day was significantly higher than on moderate pain-reinstatement day (p < 0.0001). As expected, the amount of force applied to Sham animals was greater than SNI on both moderate and high pain-reinstatement days (all p’s < 0.0001). The amount of force applied to the ipsilateral paw of SNI animals was not significantly greater on high pain-reinstatement day (p = 0.277). The von Frey test uses a logarithmic increase in force as this logarithmic increase is perceived as linear. Thus, to compare the amount of force used for stimulation for the ‘moderate’ and ‘high’ stimulation sessions between SNI and sham, we calculated the percent change in force. A Mann-Whitney test revealed that the % increase in force on ‘high’ stimulation was greater for SNI animals than sham animals (p = 0.0288; **Fig. 3c**). These analyses reveal that while the absolute amount of force delivered to SNI animals’ ipsilateral paws did not significantly differ between moderate pain-reinstatement day and high pain-reinstatement day, proportionally the increase in the amount of force per stimulation was much greater than sham animals. These analyses indicate that sham animals were stimulated with a greater amount of absolute force on high simulation reinstatement day, while SNI animals were stimulated with a greater increase of proportional force on high stimulation reinstatement day.

Importantly, while the forces used to elicit hindpaw withdrawal responses on moderate pain-reinstatement day and high pain-reinstatement day differed between SNI and Sham animals, the number of hindpaw withdrawal responses did not differ. A 2-way rmANOVA on the number of hindpaw withdrawal responses revealed a main effect of stimulation force [F(1, 20) = 0.2074, p = <0.0001; **Fig. 3d**], with greater withdrawals following ‘high’ stimulation, as expected. No main effect of surgery [F(1, 20) = 1.043, p = 0.3194] nor surgery x stimulation interaction [F(1, 20) = 0.2074, p = 0.6537] were observed, confirming successful behavioral outcome matching using these stimulation parameters.

### Chronic neuropathic pain increases sensitivity to pain-induced reinstatement

To determine the effect of painful stimulation on reinstatement of ethanol CPP, Sham and SNI mice were stimulated 10 times with either a ‘moderate’ or a ‘high’ amount of force. A one sample t-test vs. 0 revealed that moderate stimulation did not significantly increase reinstatement scores in sham mice (t = 0.7850, df =9, p = 0.4526; **Fig. 4a**), while SNI mice exhibited a significant increase (t = 3.080, df = 9, p = 0.0131). Consistent with this, SNI mice exhibited significantly greater reinstatement scores than shams (unpaired t-test; t = 2.850, df = 18, p = 0.0106; **Fig. 4a**). This indicates that animals with SNI exhibited reinstatement of ethanol seeking after moderate stimulation, while sham controls did not, suggesting that mice in chronic pain are susceptible to reinstatement of ethanol seeking in response to moderate acutely painful events.

**Figure 4.**
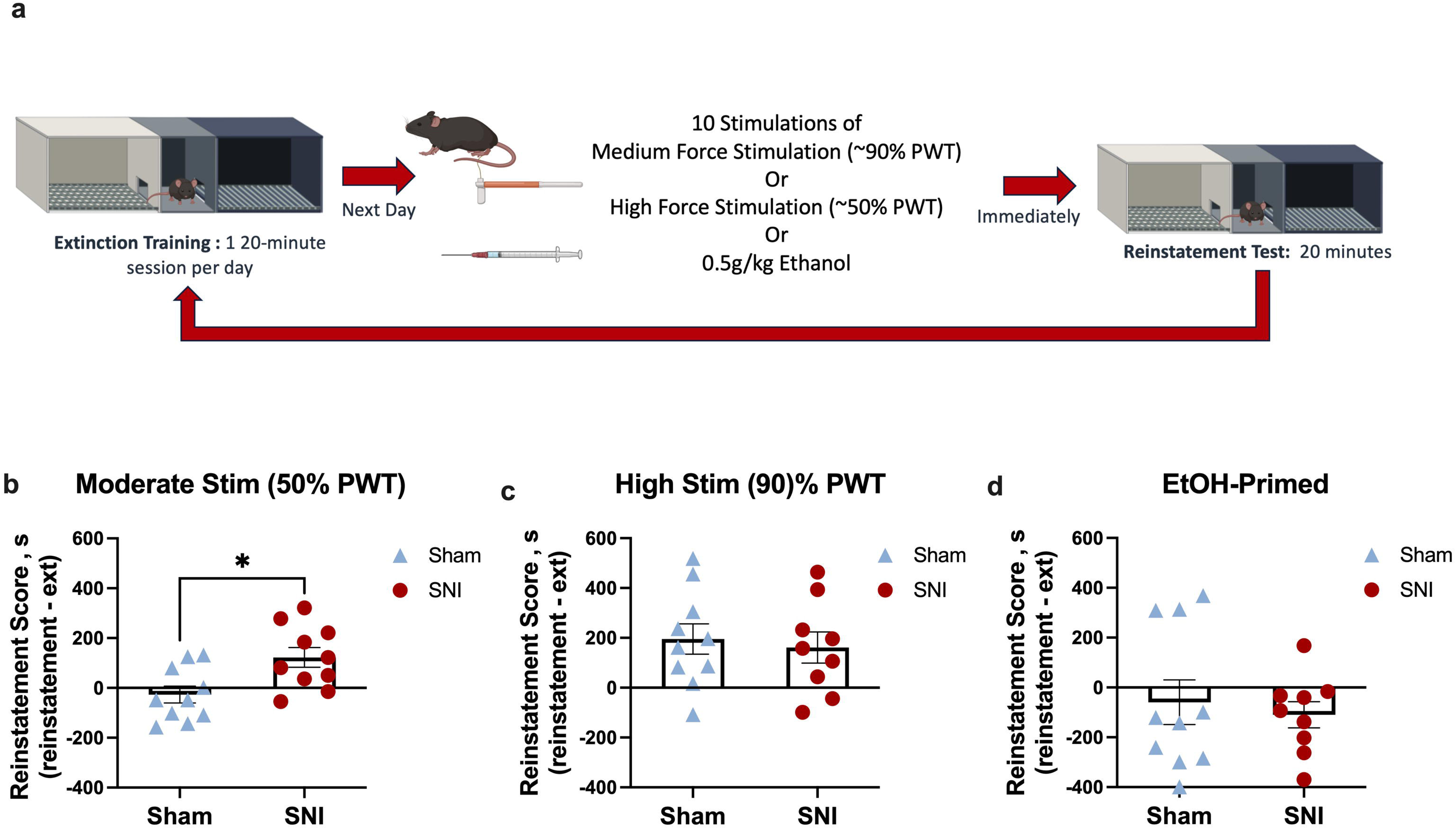
SNI increases sensitivity to pain-induced reinstatement. **(a)** Timeline of pain-induced reinstatement and ethanol-primed reinstatement. **(b)** Moderate stimulation leads to reinstatement of ethanol seeking behavior in SNI mice, but not sham mice. **(c)** High stimulation induces reinstatement of ethanol seeking of a similar magnitude in both SNI and sham mice. **(d)** Ethanol-primed reinstatement was not observed in either SNI or sham groups. Data are expressed as mean ± SEM. *p < 0.05 SNI vs. Sham

When mice experienced a ‘high’ stimulation, a one sample t-test vs. 0 indicated that both sham (t = 3.201, df =9, p = 0.0108; **Fig. 4b**) and SNI (t = 2.591, df =8, p = 0.0320) reinstated ethanol seeking. No difference was observed in the magnitude of reinstatement (unpaired t-test, t = 0.3924, df = 11, p = 0.6996). Collectively these data indicate that painful events can broadly modulate relapse-related behavior in male mice and that chronic pain increases susceptibility to this modulation.

To assess whether chronic pain modulated ethanol-primed reinstatement of ethanol CPP, a 0.5g/kg dose of ethanol was administered prior. A one sample t-test vs. 0 indicated that neither sham (t = 0.6585, df =9, p = 0.5267; **Fig. 4c**) nor SNI (t = 2.096, df =8, p = 0.0694) exhibited ethanol-primed reinstatement, though a trend was observed in SNI mice. However, reinstatement scores did not significantly differ between SNI and Sham mice (unpaired t test, t = 0.4710, df = 17, p = 0.1088), suggesting that no meaningful ethanol-primed reinstatement was observed in this paradigm.

## Discussion

Alcohol use disorder is accompanied by a host of comorbid conditions, commonly including chronic neuropathic pain (Bair et al., 2003; Von Korff et al., 2005; Zale et al., 2015) Despite evidence of a bidirectional relationship between AUD and pain, the majority of clinical and preclinical studies consider them independently. Understanding the relationship between chronic pain and AUD is difficult clinically, however, preclinical models allow for the interrogation of direct effects of chronic pain on ethanol seeking behavior (Zale et al., 2015). To expand our understanding of the interplay between chronic neuropathic pain and alcohol relapse behavior – a defining feature of AUD even after long periods of abstinence-these experiments aimed to investigate the interaction between chronic neuropathic pain and ethanol seeking/pain-induced reinstatement using the conditioned place preference assay in the spared nerve injury mouse model of chronic pain (Decosterd & Woolf, 2000). Our findings indicated that, despite similar magnitudes of ethanol CPP, male mice with chronic neuropathic exhibited elevated sensitivity to pain-induced reinstatement of ethanol seeking. This increased sensitivity was restricted to “moderate” painful stimulation, as both controls and SNI mice exhibited similar reinstatement following higher stimulation. Together, these data indicate that painful experiences can induce relapse-related behavior and those with a history of chronic neuropathic pain are more susceptible to pain-induced relapse.

In clinical populations, chronic pain is associated with relapse risk (Boissoneault et al., 2019; McDermott et al., 2018; Von Korff et al., 2005), but interrogating this relationship is challenging due to the bidirectional relationship between chronic pain and alcohol intake. While there are multiple models of relapse-related behavior in rodents, these have emphasized the ability of factors such as stress, reward-paired cues, and drug exposure to induce a return to drug seeking (Bossert et al., 2013; Kuhn et al., 2019). To our knowledge, there are no published models of pain-induced reinstatement of reward seeking for any drugs, including alcohol. Thus, we developed a model based on the conditioned place preference paradigm in which ethanol CPP was extinguished and reinstated following painful stimulation. We demonstrated that animals with and without chronic neuropathic pain (SNI and Sham mice) reinstated ethanol seeking following “high” mechanical stimulation (Fig. 4b). This suggests that intense nociceptive input can drive relapse-like behavior, and this model can be employed for studies beyond investigation of chronic neuropathic pain.

Our findings further indicated that mice with a history of chronic neuropathic pain showed greater sensitivity to reinstatement of ethanol seeking after “moderate” stimulation. Specifically, SNI – but not Sham - animals reinstated ethanol seeking after mechanical stimulations using force that causes a hindpaw withdrawal response ∼50% of the time (Fig 3C). This suggests that individuals in chronic pain are more susceptible to pain-driven relapse-like ethanol seeking. Chronic pain is associated with a dysfunction in conditioned pain modulation which is thought to occur due to dysregulation of descending modulation of inhibitory controls and can be modulated by alcohol intake(Horn-Hofmann et al., 2019). Aberrant descending modulation can lead to enhanced pain perception and motivation to seek pain relief (Gebhart, 2004; Kharkevich & Churukanov, 1999; Zhuo, 2017), including through alcohol consumption. Indeed, alcohol is thought to partially exert its analgesic effects by restoring descending pain modulation (Cutter & O’farrell, 1987; Stewart et al., 1995; Thompson et al., 2017). This chronic perturbation during chronic pain may drive selective reinstatement in SNI animals despite similar reflexive behavioral responses to “moderate” stimulation in Sham and SNI animals (**Fig. 3c**). Our findings mirror clinical data which also suggests that pain is a major cause of relapse to ethanol (Boissoneault et al., 2019).

We did not observe reinstatement following an ethanol prime in Sham or SNI mice. While others have observed ethanol-primed reinstatement (Gass & Olive, 2007; Pildervasser et al., 2014), the absence here may be due to methodological or design differences in our experiments. In particular, mice had undergone repeated pain-induced reinstatement and extinction training prior to assessment of ethanol-primed reinstatement. Although the “moderate” and “high” pain-reinstatement were counterbalanced, the inclusion of the ethanol-primed reinstatement test following the other reinstatement sessions could have resulted in a stronger extinction memory. Alternatively, the repeated painful stimulation may have altered reward seeking behavior or response to ethanol. An alternative, but not mutually exclusive, interpretation is that painful stimulation is a stronger driver of relapse-related behavior than low-dose ethanol, particularly in mice with chronic neuropathic pain. In patients with AUD, relapse is often driven by stress, negative affect, and pain rather than alcohol itself (Boissoneault et al., 2019; Walitzer & Dearing, 2006). Thus, reducing pain and negative affective states may suppress relapse-risk.

This study used a dose of ethanol (0.5 g/kg) for CPP that we previously demonstrated was able to reverse both mechanical allodynia and altered dynamic weight bearing in SNI male mice (Nothem et al., 2023). Those findings were replicated here as we again demonstrated that in SNI mice, paw withdrawal thresholds were reduced after SNI surgery, and this was reversed by 0.5g/kg ethanol injection. No effect was observed in sham mice (**Fig 1b**), consistent with our previous findings demonstrating antiallodynic effects on evoked mechanical stimulation in SNI mice without producing antinociceptive effects in sham animals. A similar pattern was observed in measures of hindpaw weight bearing—a non-reflexive proxy for ongoing pain. SNI mice exhibited reduced weight bearing on the injured hindpaw after surgery, which was again reversed by 0.5g/kg ethanol (**Fig. 1c**). As we reported previously, the 0.5g/kg of ethanol did not produce reductions in movement during CPP conditioning (**Fig. 2a**), suggesting that this dose was not sedating which would confound our interpretation of the results from the pain behavioral assays.

The 0.5g/kg dose is a relatively low dose of ethanol and does not consistently yield a CPP in male mice (Bozarth, 1990; Groblewski et al., 2008; Shimizu et al., 1935). Thus, we initially hypothesized that we would observe a CPP only in SNI, but not sham, mice, potentially through the anti-allodynic/negative reinforcing properties of this dose of ethanol. However, Sham and SNI mice formed an ethanol CPP that was similar in magnitude, consistent with similar reward-context conditioning. Additionally, the magnitude ethanol CPP we observed was similar to what others have reported, even when using higher doses (Groblewski et al., 2008; Shimizu et al., 1935). It is possible that reducing the number of pairing sessions may have yielded differences in CPP magnitude between sham and SNI animals. Consistent with this, others have found that persistent pain facilitated morphine CPP, reflected by a CPP for a lower dose and requiring fewer conditioning sessions (Cahill et al., 2013; Zhang et al., 2014).

Previous research has highlighted the bidirectional relationship between chronic pain and alcohol use, with studies in both humans and animal models showing that pain can drive increased ethanol consumption. Voluntary ethanol consumption is increased in male mice undergoing chronic pain (Butler et al., 2017; Trouvin et al., 2022; Yin et al., 2024; Yu et al., 2019) and rodents in chronic pain will readily seek and self-administer other analgesics (Hipólito et al., 2015). While in our hands SNI does not appear to enhance ethanol CPP under the current conditions, chronic pain may still drive increased ethanol seeking and intake through mechanisms independent of conditioned ethanol reward. Alcohol use in the context of chronic pain likely involves both negative and positive reinforcement processes. Negative reinforcement occurs when an aversive state such as pain - characterized by both sensory discomfort and emotional distress - drives behavior aimed at alleviating that state (Baker et al., 2004). For individuals with chronic pain, successfully obtaining analgesia by consumption of alcohol or other analgesics can be negatively reinforcing due to relief from the persistent aversive state (Egli et al., 2012; Koob & Volkow, 2010; Volkow et al., 2016). This is especially relevant during tonic pain states, where the ongoing aversive nature of pain exacerbates the motivation to seek relief. In contrast, positive reinforcement is mediated by alcohol’s rewarding properties including its capacity to induce euphoria and relaxation (Stewart et al., 1995; Thompson et al., 2017). These properties are not mutually exclusive and may interact in complex ways in the context of chronic pain (De Aquino et al., 2024). It is plausible that sham animals formed a place preference due to the positive rewarding effects of ethanol, while SNI animals, which are likely experiencing a chronic aversive state, exhibited ethanol seeking due to negative reinforcement. However, the CPP task as employed here did not enable this dissociation which is an exciting area for future study.

### Caveats and Future Directions

The current findings provide insight into the relationship between chronic pain and ethanol-seeking behavior, particularly in the context of relapse vulnerability. Several key questions remained to be answered. While we were able to show that painful stimulation can induce relapse-related behavior using a model of CPP, future studies should investigate whether painful stimulation can increase ethanol seeking in an operant self-administration setting. Furthermore, while we used a mechanical stimulus to induce pain, we do not know if the increase in ethanol seeking is specific to that modality and so future studies should investigate whether thermal or chemical stimuli can produce same effect on relapse-related behaviors. It is also possible that pain may potentiate other drivers of reinstatement, such as drug-paired cues or stress. Acutely painful events cause stress and release cortisol (Muhtz et al., 2013), and while this study did not delineate between the pain experience and stress mechanisms future studies should delineate the role that pain-induced stress plays in ethanol seeking behaviors. Given the inability to disentangle the contribution of positive versus negative reinforcement to the development, expression, and reinstatement of ethanol CPP, future studies involving operant self-administration and characterizing the contributions of discrete neural substrates/circuits will provide insight into the processes governing relapse-related behavior in individuals with chronic pain.

Importantly, in this study, only male mice were used. This was due to sex differences in the analgesic properties of ethanol in mice, as we have previously reported that female mice exhibited limited reversal of pain-related behaviors in both the von Frey and dynamic weight bearing assays (Nothem et al., 2023). While most patients with AUD are males (Dufour, 1995; Erol & Karpyak, 2015; Khan et al., 2013), this gap is closing (Keyes et al., 2008; Slade et al., 2016; White, 2020). Sex differences in ethanol CPP(Barros-Santos et al., 2021; Cunningham & Shields, 2018; Xie et al., 2019), operant self-administration(Priddy et al., 2017a; Randall et al., 2017) and home-cage (Priddy et al., 2017b; Rivera-Irizarry et al., 2023) consumption are frequently observed in rodent models. Further, sex differences are also reported in chronic neuropathic pain in clinical populations (Marcianò et al., 2024; Miclescu et al., 2022) and rodent models (Boullon et al., 2021; Gregus et al., 2021; Lee et al., 2023). Thus, it will be essential to characterize the interaction between chronic pain and AUD in women and female preclinical models.

## Conclusions

Despite a growing body of evidence of frequent comorbidity between chronic pain and alcohol use disorder, significant gaps remain in understanding the behavioral processes underlying ethanol use in chronic pain conditions. Expansion of preclinical models of pain-driven relapse will enable interrogation of the underlying neural and behavioral mechanisms that lead to increased relapse vulnerability in individuals experiencing pain. Our findings using a novel model of pain-driven relapse-related behavior indicate that pain-induced reinstatement of ethanol seeking is increased following a history of chronic neuropathic pain. There remains a lack of effective therapeutic approaches for the treatment of AUD, which may reflect the paucity of complex models of AUD that capture common comorbidities. Given the large patient population of individuals with comorbid chronic pain and AUD, future studies on AUD should consider pain or chronic pain as an important variable. The use of this model of pain-induced reinstatement to identify neurobehavioral substrates of relapse during chronic pain may thus inform strategies to attain and maintain abstinence in people living with AUD.

## Notes

Funding sources: This research was supported by the National Institute on Alcohol Abuse and Alcoholism under Award Number R21AA027629 (JMB).

### Competing Interest Statement

The authors have declared no competing interest.

## References

Alford, D. P., German, J. S., Samet, J. H., Cheng, D. M., Lloyd-Travaglini, C. A., & Saitz, R. (2016). Primary Care Patients with Drug Use Report Chronic Pain and Self-Medicate with Alcohol and Other Drugs. Journal of General Internal Medicine, 31(5), 486–491. 10.1007/s11606-016-3586-5

Andrew, R., Derry, S., Taylor, R. S., Straube, S., Phillips, C. J., Moore, R. A., Derry, S., Taylor, R. S., Straube, S., & Phillips, C. J. (2014). The costs and consequences of adequately managed chronic non-cancer pain and chronic neuropathic pain. Pain Practice, 14(1), 79–94. 10.1111/papr.12050

Axley, P. D., Richardson, C. T., & Singal, A. K. (2019). Epidemiology of Alcohol Consumption and Societal Burden of Alcoholism and Alcoholic Liver Disease. In Clinics in Liver Disease (Vol. 23, Issue 1, pp. 39–50). W.B. Saunders. 10.1016/j.cld.2018.09.011

Bair, M. J., Robinson, R. L., Katon, W., & Kroenke, K. (2003). Depression and Pain Comorbidity. Archives of Internal Medicine. 10.1001/archinte.163.20.2433

Baker, T. B., Piper, M. E., McCarthy, D. E., Majeskie, M. R., & Fiore, M. C. (2004). Addiction Motivation Reformulated: An Affective Processing Model of Negative Reinforcement. Psychological Review, 111(1), 33–51. 10.1037/0033-295X.111.1.33

Barros-Santos, T., Libarino-Santos, M., Anjos-Santos, A., Lins, J. F., Leite, J. P. C., Pacheco, R. C., Nascimento-Rocha, V., Kisaki, N. D., Tamura, E. K., Oliveira-Lima, A. J., Berro, L. F., Uetanabaro, A. P. T., Nicoli, J. R., & Marinho, E. A. V. (2021). Sex differences in the development of conditioned place preference induced by intragastric alcohol administration in mice. Drug and Alcohol Dependence, 229. 10.1016/j.drugalcdep.2021.109105

Boissoneault, J., Lewis, B., & Nixon, S. J. (2019). Characterizing chronic pain and alcohol use trajectory among treatment-seeking alcoholics. Alcohol, 75. 10.1016/j.alcohol.2018.05.009

Bossert, J. M., Marchant, N. J., Calu, D. J., & Shaham, Y. (2013). The reinstatement model of drug relapse: Recent neurobiological findings, emerging research topics, and translational research. In Psychopharmacology (Vol. 229, Issue 3, pp. 453–476). 10.1007/s00213-013-3120-y

Boullon, L., Finn, D. P., & Llorente-Berzal, Á. (2021). Sex Differences in a Rat Model of Peripheral Neuropathic Pain and Associated Levels of Endogenous Cannabinoid Ligands. Frontiers in Pain Research, 2. 10.3389/fpain.2021.673638

Bozarth, M. A. (1990). Evidence for the rewarding effects of ethanol using the conditioned place preference method. Pharmacology, Biochemistry, and Behavior, 35(2), 485–487. 10.1016/0091-3057(90)90191-j

Brennan, P. L., Schutte, K. K., & Moos, R. H. (2005). Pain and use of alcohol to manage pain: prevalence and 3-year outcomes among older problem and non-problem drinkers. Addiction (Abingdon, England), 100(6), 777–786. 10.1111/j.1360-0443.2005.01074.x

Butler, R. K., Knapp, D. J., Ulici, V., Longobardi, L., Loeser, R. F., & Breese, G. R. (2017). A mouse model for chronic pain-induced increase in ethanol consumption. Pain, 158(3), 457–462. 10.1097/j.pain.0000000000000780

Cahill, C. M., Xue, L., Grenier, P., Magnussen, C., Lecour, S., & Olmstead, M. C. (2013). Changes in morphine reward in a model of neuropathic pain. Behavioural Pharmacology, 24(3), 207–213. 10.1097/FBP.0b013e3283618ac8

Chaplan, S. R., Bach, F. W., Pogrel, J. W., Chung, J. M., & Yaksh, T. L. (1994). Quantitative assessment of tactile allodynia in the rat paw. In Journal of Neuroscience Methods (Vol. 53).

Cunningham, C. L., & Shields, C. N. (2018). Effects of sex on ethanol conditioned place preference, activity and variability in C57BL/6J and DBA/2J mice. Pharmacology Biochemistry and Behavior, 173, 84–89. 10.1016/j.pbb.2018.07.008

Cutter, H. S. G., & O’farrell, T. J. (1987). EXPERIENCE WITH ALCOHOL AND THE ENDOGENOUS OPIOID SYSTEM IN ETHANOL ANALGESIA. In Addictive Behaviors (Vol. 12).

De Aquino, J. P., Sloan, M. E., Nunes, J. C., Costa, G. P. A., Katz, J. L., de Oliveira, D., Ra, J., Tang, V. M., & Petrakis, I. L. (2024). Alcohol Use Disorder and Chronic Pain: An Overlooked Epidemic. In The American journal of psychiatry (Vol. 181, Issue 5, pp. 391– 402). 10.1176/appi.ajp.20230886

Decosterd, I., & Woolf, C. J. (2000). Spared nerve injury: An animal model of persistent peripheral neuropathic pain. Pain. 10.1016/S0304-3959(00)00276-1

Dixon, W. J. (1980). EFFICIENT ANALYSIS OF EXPERIMENTAL OBSERVATIONS. In Ann. Rev. Pharmacol. Toxicol. 198U (Vol. 20). www.annualreviews.org

Dufour, M. C. (1995). Twenty-Five Years of Alcohol Epidemiology Trends, Techniques, and Transitions.

Dundee, J. W., Martin Isaac, F. F. A. R. C. S., Richard, D. A., Clarke, S. J., Belfast, F. F. A. R. C. S., & Ireland, N. (1969). Use of Alcohol in Anesthesia. http://journals.lww.com/anesthesia-analgesia

Egli, M., Koob, G. F., & Edwards, S. (2012). Alcohol dependence as a chronic pain disorder. Neuroscience and Biobehavioral Reviews, 36(10), 2179–2192. 10.1016/j.neubiorev.2012.07.010

Erichsen, H. K., & Blackburn-Munro, G. (2002). Pharmacological characterisation of the spared nerve injury model of neuropathic pain. Pain, 98(1–2), 151–161. S0304395902000398

Erol, A., & Karpyak, V. M. (2015). Sex and gender-related differences in alcohol use and its consequences: Contemporary knowledge and future research considerations. In Drug and Alcohol Dependence (Vol. 156, pp. 1–13). Elsevier Ireland Ltd. 10.1016/j.drugalcdep.2015.08.023

Gass, J. T., & Olive, M. F. (2007). Reinstatement of ethanol-seeking behavior following intravenous self-administration in Wistar rats. Alcoholism, Clinical and Experimental Research, 31(9), 1441–1445. 10.1111/j.1530-0277.2007.00480.x

Gebhart, G. F. (2004). Descending modulation of pain. 27, 729–737. 10.1016/j.neubiorev.2003.11.008

Gregus, A. M., Levine, I. S., Eddinger, K. A., Yaksh, T. L., & Buczynski, M. W. (2021). Sex differences in neuroimmune and glial mechanisms of pain. In Pain (Vol. 162, Issue 8, pp. 2186–2200). Lippincott Williams and Wilkins. 10.1097/j.pain.0000000000002215

Groblewski, P. A., Bax, L. S., & Cunningham, C. L. (2008). Reference-dose place conditioning with ethanol in mice: Empirical and theoretical analysis. Psychopharmacology, 201(1), 97–106. 10.1007/s00213-008-1251-3

Guida, F., De Gregorio, D., Palazzo, E., Ricciardi, F., Boccella, S., Belardo, C., Iannotta, M., Infantino, R., Formato, F., Marabese, I., Luongo, L., de Novellis, V., & Maione, S. (2020). Behavioral, Biochemical and Electrophysiological Changes in Spared Nerve Injury Model of Neuropathic Pain. International Journal of Molecular Sciences, 21(9), 1–21. 10.3390/ijms21093396

Hipólito, L., Wilson-Poe, A., Campos-Jurado, Y., Zhong, E., Gonzalez-Romero, J., Virag, L., Whittington, R., Comer, S. D., Carlton, S. M., Walker, B. M., Bruchas, M. R., & Morón, J. A. (2015). Inflammatory pain promotes increased opioid self-administration: Role of dysregulated ventral tegmental area µ opioid receptors. Journal of Neuroscience, 35(35). 10.1523/JNEUROSCI.1053-15.2015

Horn-Hofmann, C., Büscher, P., Lautenbacher, S., & Wolstein, J. (2015). The effect of nonrecurring alcohol administration on pain perception in humans: A systematic review. Journal of Pain Research, 8, 175–187. 10.2147/JPR.S79618

Horn-Hofmann, C., Capito, E. S., Wolstein, J. O., & Lautenbachera, S. (2019). Acute alcohol effects on conditioned pain modulation, but not temporal summation of pain. Pain, 160(9), 2063–2071. 10.1097/j.pain.0000000000001597

Keyes, K. M., Grant, B. F., & Hasin, D. S. (2008). Evidence for a closing gender gap in alcohol use, abuse, and dependence in the United States population. Drug and Alcohol Dependence, 93(1–2), 21–29. 10.1016/j.drugalcdep.2007.08.017

Khan, S., Okuda, M., Hasin, D. S., Secades-Villa, R., Keyes, K., Lin, K. H., Grant, B., & Blanco, C. (2013). Gender differences in lifetime alcohol dependence: Results from the national epidemiologic survey on alcohol and related conditions. Alcoholism: Clinical and Experimental Research, 37(10), 1696–1705. 10.1111/acer.12158

Kharkevich, D. A., & Churukanov, V. V. (1999). Pharmacological regulation of descending cortical control of the nociceptive processing. European Journal of Pharmacology, 375(1– 3), 121–131. 10.1016/S0014-2999(99)00264-2

King, C. D., Goodin, B., Kindler, L. L., Caudle, R. M., Edwards, R. R., Gravenstein, N., Riley, J. L., & Fillingim, R. B. (2013). Reduction of conditioned pain modulation in humans by naltrexone: An exploratory study of the effects of pain catastrophizing. Journal of Behavioral Medicine, 36(3), 315–327. 10.1007/s10865-012-9424-2

Koob, G. F., & Volkow, N. D. (2010). Neurocircuitry of addiction. Neuropsychopharmacology, 35(1), 217–238. 10.1038/npp.2009.110

Kuhn, B. N., Kalivas, P. W., & Bobadilla, A. C. (2019). Understanding Addiction Using Animal Models. In Frontiers in Behavioral Neuroscience (Vol. 13). Frontiers Media S.A. 10.3389/fnbeh.2019.00262

Lee, S. E., Greenough, E. K., Oancea, P., Scheinfeld, A. R., Douglas, A. M., & Gaudet, A. D. (2023). Sex Differences in Pain: Spinal Cord Injury in Female and Male Mice Elicits Behaviors Related to Neuropathic Pain. Journal of Neurotrauma, 40(9–10), 833–844. 10.1089/neu.2022.0482

Marcianò, G., Siniscalchi, A., Di Gennaro, G., Rania, V., Vocca, C., Palleria, C., Catarisano, L., Muraca, L., Citraro, R., Evangelista, M., De Sarro, G., D’Agostino, B., Abrego-Guandique, D. M., Cione, E., Morlion, B., & Gallelli, L. (2024). Assessing Gender Differences in Neuropathic Pain Management: Findings from a Real-Life Clinical Cross-Sectional Observational Study. Journal of Clinical Medicine, 13(19). 10.3390/jcm13195682

McDermott, K. A., Joyner, K. J., Hakes, J. K., Okey, S. A., & Cougle, J. R. (2018). Pain interference and alcohol, nicotine, and cannabis use disorder in a national sample of substance users. Drug and Alcohol Dependence, 186, 53–59. 10.1016/j.drugalcdep.2018.01.011

Miclescu, A. A., Gkatziani, P., Granlund, P., Butler, S., & Gordh, T. (2022). Sex-related differences in experimental pain sensitivity in subjects with painful or painless neuropathy after surgical repair of traumatic nerve injuries. Pain Reports, 7(6), E1033. 10.1097/PR9.0000000000001033

Mogil, J. S., Graham, A. C., Ritchie, J., Hughes, S. F., Austin, J. S., Schorscher-Petcu, A., Langford, D. J., & Bennett, G. J. (2010). Hypolocomotion, asymmetrically directed behaviors (licking, lifting, flinching, and shaking) and dynamic weight bearing (gait) changes are not measures of neuropathic pain in mice. Molecular Pain, 6. 10.1186/1744-8069-6-34

Muhtz, C., Rodriguez-Raecke, R., Hinkelmann, K., Moeller-Bertram, T., Kiefer, F., Wiedemann, K., May, A., & Otte, C. (2013). Cortisol response to experimental pain in patients with chronic low back pain and patients with major depression. *Pain Medicine (Malden*, Mass.), 14(4), 498–503. 10.1111/j.1526-4637.2012.01514.x

Niesters, M., Proto, P. L., Aarts, L., Sarton, E. Y., Drewes, A. M., & Dahan, A. (2014). Tapentadol potentiates descending pain inhibition in chronic pain patients with diabetic polyneuropathy. British Journal of Anaesthesia, 113(1), 148–156. 10.1093/bja/aeu056

Nothem, M. A., Wickman, J. R., Giacometti, L. L., & Barker, J. M. (2023). Effects of ethanol on mechanical allodynia and dynamic weight bearing in male and female mice with spared nerve injury. Alcoholism: Clinical and Experimental Research, 47(2). 10.1111/acer.14997

Perrino, A. C., Ralevski, E., Acampora, G., Edgecombe, J., Limoncelli, D., & Petrakis, I. L. (2008). Ethanol and pain sensitivity: Effects in healthy subjects using an acute pain paradigm. Alcoholism: Clinical and Experimental Research, 32(6), 952–958. 10.1111/j.1530-0277.2008.00653.x

Pildervasser, J. V. N., Abrahao, K. P., & Souza-Formigoni, M. L. O. (2014). Distinct behavioral phenotypes in ethanol-induced place preference are associated with different extinction and reinstatement but not behavioral sensitization responses. Frontiers in Behavioral Neuroscience, 8(AUG). 10.3389/fnbeh.2014.00267

Priddy, B. M., Carmack, S. A., Thomas, L. C., Vendruscolo, J. C. M., Koob, G. F., & Vendruscolo, L. F. (2017a). Sex, strain, and estrous cycle influences on alcohol drinking in rats. Pharmacology Biochemistry and Behavior, 152, 61–67. 10.1016/j.pbb.2016.08.001

Priddy, B. M., Carmack, S. A., Thomas, L. C., Vendruscolo, J. C. M., Koob, G. F., & Vendruscolo, L. F. (2017b). Sex, strain, and estrous cycle influences on alcohol drinking in rats. Pharmacology Biochemistry and Behavior, 152, 61–67. 10.1016/j.pbb.2016.08.001

Randall, P. A., Stewart, R. T., & Besheer, J. (2017). Sex differences in alcohol self-administration and relapse-like behavior in Long-Evans rats. Pharmacology Biochemistry and Behavior, 156, 1–9. 10.1016/j.pbb.2017.03.005

Rivera-Irizarry, J. K., Zallar, L. J., Levine, O. B., Skelly, M. J., Boyce, J. E., Barney, T., Kopyto, R., & Pleil, K. E. (2023). Sex differences in binge alcohol drinking and the behavioral consequences of protracted abstinence in C57BL/6J mice. Biology of Sex Differences, 14(1). 10.1186/s13293-023-00565-0

Shimizu, C., Oki, Y., Mitani, Y., Nakamura, T., & Nabeshima, T. (1935).) and c NPO Japanese Drug Organization of Appropriate Use and Research. In Biol. Pharm. Bull (Vol. 38, Issue 12).

Slade, T., Chapman, C., Swift, W., Keyes, K., Tonks, Z., & Teesson, M. (2016). Birth cohort trends in the global epidemiology of alcohol use and alcohol-related harms in men and women: systematic review and meta regression. BMJ Open, 6, 11827. 10.1136/bmjopen-2016

Stewart, S. H., Finn, P. R., Pihl, R. O., Stewart, S. H., Finn, P. R., & Pihl, R. O. (1995). A dose-response study of the effects of alcohol on the perceptions of pain and discomfort due to electric shock in men at high familial-genetic risk for alcoholism. In Psychopharmacology (Vol. 119).

Thompson, T., Oram, C., Correll, C. U., Tsermentseli, S., & Stubbs, B. (2017). Analgesic Effects of Alcohol: A Systematic Review and Meta-Analysis of Controlled Experimental Studies in Healthy Participants. In Journal of Pain (Vol. 18, Issue 5, pp. 499–510). Churchill Livingstone Inc. 10.1016/j.jpain.2016.11.009

Trouvin, A. P., Attal, N., & Perrot, S. (2022). Association between alcohol consumption and pain a systematic review and meta-analysis. In British Journal of Anaesthesia (Vol. 129, Issue 3, pp. 278–281). Elsevier Ltd. 10.1016/j.bja.2022.06.006

Volkow, N. D., Koob, G. F., & McLellan, A. T. (2016). Neurobiologic advances from the brain disease model of addiction. New England Journal of Medicine, 374(4), 363–371. 10.1056/NEJMra1511480

Von Korff, M., Crane, P., Lane, M., Miglioretti, D. L., Simon, G., Saunders, K., Stang, P., Brandenburg, N., & Kessler, R. (2005). Chronic spinal pain and physical-mental comorbidity in the United States: Results from the national comorbidity survey replication. Pain, 113(3), 331–339. 10.1016/j.pain.2004.11.010

Walitzer, K. S., & Dearing, R. L. (2006). Gender differences in alcohol and substance use relapse. Clinical Psychology Review, 26(2), 128–148. 10.1016/j.cpr.2005.11.003

White, A. M. (2020). Gender differences in the epidemiology of alcohol use and related harms in the United States. Alcohol Research: Current Reviews, 40(2). 10.35946/arcr.v40.2.01

Wolff, H. G., Hardy, J. D., & Goodell, H. (1940). MEASUREMENT OF THE EFFECT ON THE PAIN THRESHOLD OF ACETYLSALICYLIC ACID.

Woodrow, K. M., & Eltherington, L. G. (1988). Feeling no pain: alcohol as an analgesic. In Pain (Vol. 32).

Xie, Q., Buck, L. A., Bryant, K. G., & Barker, J. M. (2019). Sex Differences in Ethanol Reward Seeking Under Conflict in Mice. Alcoholism: Clinical and Experimental Research, 43(7), 1556–1566. 10.1111/acer.14070

Yin, Y., Haggerty, D. L., Zhou, S., Atwood, B. K., & Sheets, P. L. (2024). Converging Effects of Chronic Pain and Binge Alcohol Consumption on Anterior Insular Cortex Neurons Projecting to the Dorsolateral Striatum in Male Mice. Journal of Neuroscience, 44(16). 10.1523/JNEUROSCI.1287-23.2024

Yu, W., Hwa, L. S., Makhijani, V. H., Besheer, J., & Kash, T. L. (2019). Chronic inflammatory pain drives alcohol drinking in a sex-dependent manner for C57BL/6J mice. Alcohol (Fayetteville, N.Y.), 77, 135. 10.1016/J.ALCOHOL.2018.10.002

Zale, E. L., Maisto, S. A., & Ditre, J. W. (2015). Interrelations between pain and alcohol: An integrative review. In Clinical Psychology Review (Vol. 37, pp. 57–71). Elsevier Inc. 10.1016/j.cpr.2015.02.005

Zhang, Z., Tao, W., Hou, Y. Y., Wang, W., Lu, Y. G., & Pan, Z. Z. (2014). Persistent pain facilitates response to morphine reward by downregulation of central amygdala GABAergic function. Neuropsychopharmacology, 39(9), 2263–2271. 10.1038/npp.2014.77

Zhuo, M. (2017). Descending facilitation: From basic science to the treatment of chronic pain. Molecular Pain, 13, 1–12. 10.1177/1744806917699212

